# A type III CRISPR ancillary ribonuclease degrades its cyclic oligoadenylate activator

**DOI:** 10.1101/582114

**Authors:** Januka S. Athukoralage, Shirley Graham, Sabine Grüschow, Christophe Rouillon, Malcolm F. White

**Author notes:** Present address: Max Planck Unit for the Science of Pathogens, Charitéplatz 1, 10117 Berlin, Germany (Christophe Rouillon).

## Abstract

Cyclic oligoadenylate (cOA) secondary messengers are generated by type III CRISPR systems in response to viral infection. cOA allosterically activates the CRISPR ancillary ribonucleases Csx1/Csm6, which degrade RNA non-specifically using a HEPN (Higher Eukaryotes and Prokaryotes, Nucleotide binding) active site. This provides effective immunity, but can also lead to growth arrest in infected cells, necessitating a means to deactivate the ribonuclease once viral infection has been cleared. In the crenarchaea, dedicated ring nucleases degrade cA_4_ (cOA consisting of 4 AMP units), but the equivalent enzyme has not been identified in bacteria. We demonstrate that, in *Thermus thermophilus* HB8, the uncharacterised protein TTHB144 is a cA_4_-activated HEPN ribonuclease that also degrades its activator. TTHB144 binds and degrades cA_4_ at an N-terminal CARF (CRISPR Associated Rossman Fold) domain. The two activities can be separated by site-directed mutagenesis. TTHB144 is thus the first example of a self-limiting CRISPR ribonuclease.

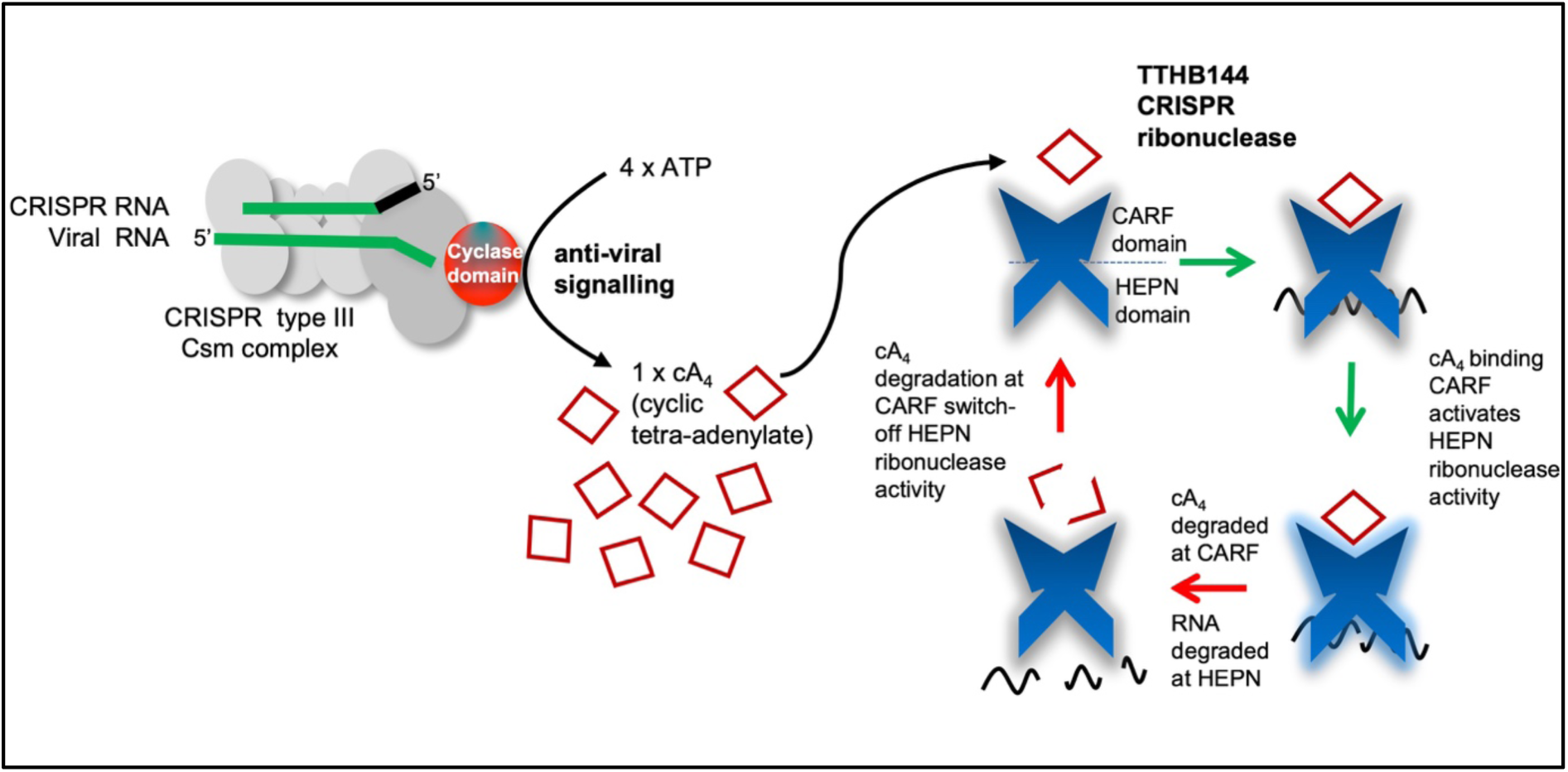
Graphical abstract.

## Introduction

The CRISPR system provides prokaryotes with adaptive immunity against mobile genetic elements (MGEs; reviewed in [1–3]). Type III (Csm/Cmr) CRISPR effector complexes harbour two nuclease activities for defence against MGEs: cleavage of foreign “target” RNA by the Cas7 subunit and degradation of single-stranded DNA by the HD nuclease domain (reviewed in [4,5]). Additionally, effector complexes produce cyclic oligoadenylate (cOA) anti-viral signalling molecules that activate CRISPR ancillary proteins to potentiate the immune response [6,7]. On target RNA binding, the cyclase domain of the Cas10 subunit polymerises ATP into cOA anti-viral signalling molecules, which consist of 3-6, 5′ to 3′ linked AMP subunits [6–9]. cOA acts as an ‘alarm signal’ within cells and strongly stimulates the activity of the CRISPR ancillary ribonucleases Csx1 and Csm6 [6–8]. Csx1/Csm6 consist of a CARF (CRISPR Associated Rossman Fold) domain that binds cOA and a HEPN (Higher Eukaryotes and Prokaryotes Nucleotide binding) domain that possess weak ribonuclease activity in the absence of cOA [10,11]. Once stimulated by cOA, the non-specific RNA degradation activity of these ribonucleases impacts both viral and cell growth [12]. Therefore, to recover from viral infection, cells require a mechanism for the removal of cOA. *Sulfolobus solfataricus (Sso)* encodes dedicated ring nucleases which degrade the cyclic tetra-adenylate (cA_4_) activator and deactivate Csx1 [13]. Thus far, ring nucleases have only been identified in the crenarchaea, and, as highlighted by Mo and Marraffini [14], the enzyme(s) responsible for cOA degradation in bacteria remain unknown. The type III CRISPR system of the bacterium *Thermus thermophilus* HB8 has been investigated [15,16], and its CRISPR ancillary ribonuclease TTHB152 was among the first shown to be activated by cA_4_ [10]. The type III CRISPR locus of *T. thermophilus* also encodes an uncharacterised CARF domain-containing protein, TTHB144, which was reported to be Csm6-like [17]. Here we report that TTHB144 is also a potent CRISPR ancillary ribonuclease activated by cA_4_, and degrades cA_4_ using its CARF domain. This enzyme therefore represents the first known example of a cOA dependent enzyme that degrades its own activator.

## Results and Discussion

The *Thermus thermophilus* HB8 type III CRISPR locus encodes three CARF domain containing proteins, TTHB144, 152 and 155 (Figure 1a). A X-ray crystal structure is available for TTHB152 (PDB: 5FSH) revealing a dimeric protein consisting of N-terminal CARF and C-terminal HEPN domain [10]. We modelled the structure of TTHB144 using the Phyre^2^ server [18], using TTHB152 as a template, and modelled cA_4_ into the electropositive pocket within the dimeric CARF domain (Figure 1b&c). Multiple sequence alignment identified highly conserved arginine and histidine residues within the HEPN domain characteristic of the Rx4-6H motif of HEPN nucleases [19]. Furthermore, we observed conserved lysine (K94) and threonine (T10/T11) residues within the ligand binding pocket of the CARF domain. By analogy with the ring nuclease Sso2081 [13], residues K94 and T10/T11 are suitably positioned to interact with the cA_4_ ligand. Consequently, we constructed a synthetic gene encoding TTHB144, expressed the protein in *E. coli* using the plasmid pEV5hisTEV [20] and purified the recombinant protein using immobilised metal affinity and size exclusion chromatography [21], allowing investigation of its enzymatic activities. Site directed protein variants were constructed and purified as for the wild-type enzyme.

**Figure 1.**
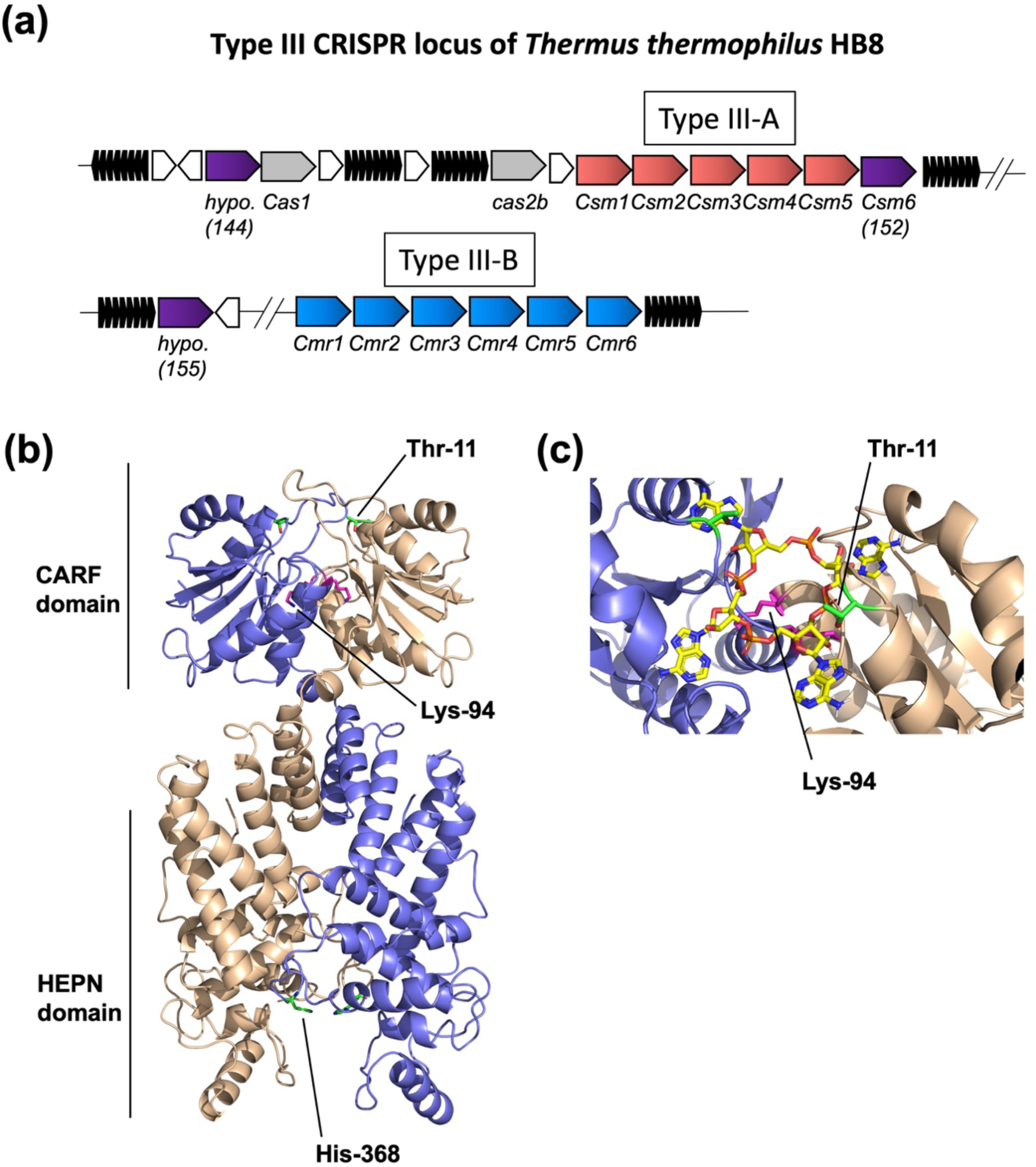
Type III CRISPR locus of *Thermus thermophilus (TTHB)* HB8 and predicted structure of TTHB144 with cA_4_ modelled into the CARF domain. (**a**) Gene neighbourhood of *TTHB144* encoded on plasmid pTT27. Three genes encoding CARF domain-containing proteins (shown in purple) are present in the type III CRISPR locus of TTHB. TTHB152 is a Csm6 family protein, while TTHB144 and TTHB155 are hypothetical proteins of unknown function. (**b**) TTHB144 structure modelled using Phyre2. Each subunit of the predicted homodimer is shown by a different colour (blue or cream). The highly conserved residues Thr-11, Lys-94 and His-368 are shown. (**c**) cA_4_ modelled into the CARF domain of TTHB144. Lys-94 is situated centrally beneath the cA_4_ molecule, and the side-chain of Thr-11 is suitably positioned to participate in catalysis of cA_4_ cleavage.

TTHB144 exhibited potent ribonuclease activity in the presence of cA_4_ and degraded RNA non-specifically (Figure 2a). The H368A variant, targeting the HEPN active site, had no RNase activity, confirming that TTHB144 is a canonical HEPN ribonuclease. The T10A/T11A variant was still an active cA_4_-dependent ribonuclease, confirming cA_4_ binding in this variant, but the K94A variant was inactive, suggesting that cA_4_ no longer binds to activate the ribonuclease. Using fluorimetry and the RNaseAlert(tm) assay system (Integrated DNA Technologies, USA) [6], we followed cA_4_ activated RNA cleavage by TTHB144 in a continuous assay (Figure 2b). Consistent with observations made for other CRISPR ancillary ribonucleases such as *Enterococcus italicus* Csm6, *Streptococcus thermophilus* Csm6 and TTHB152 [10], fluorimetry revealed weak TTHB144 ribonuclease activity which was greatly enhanced by the addition of cA_4_.

**Figure 2.**
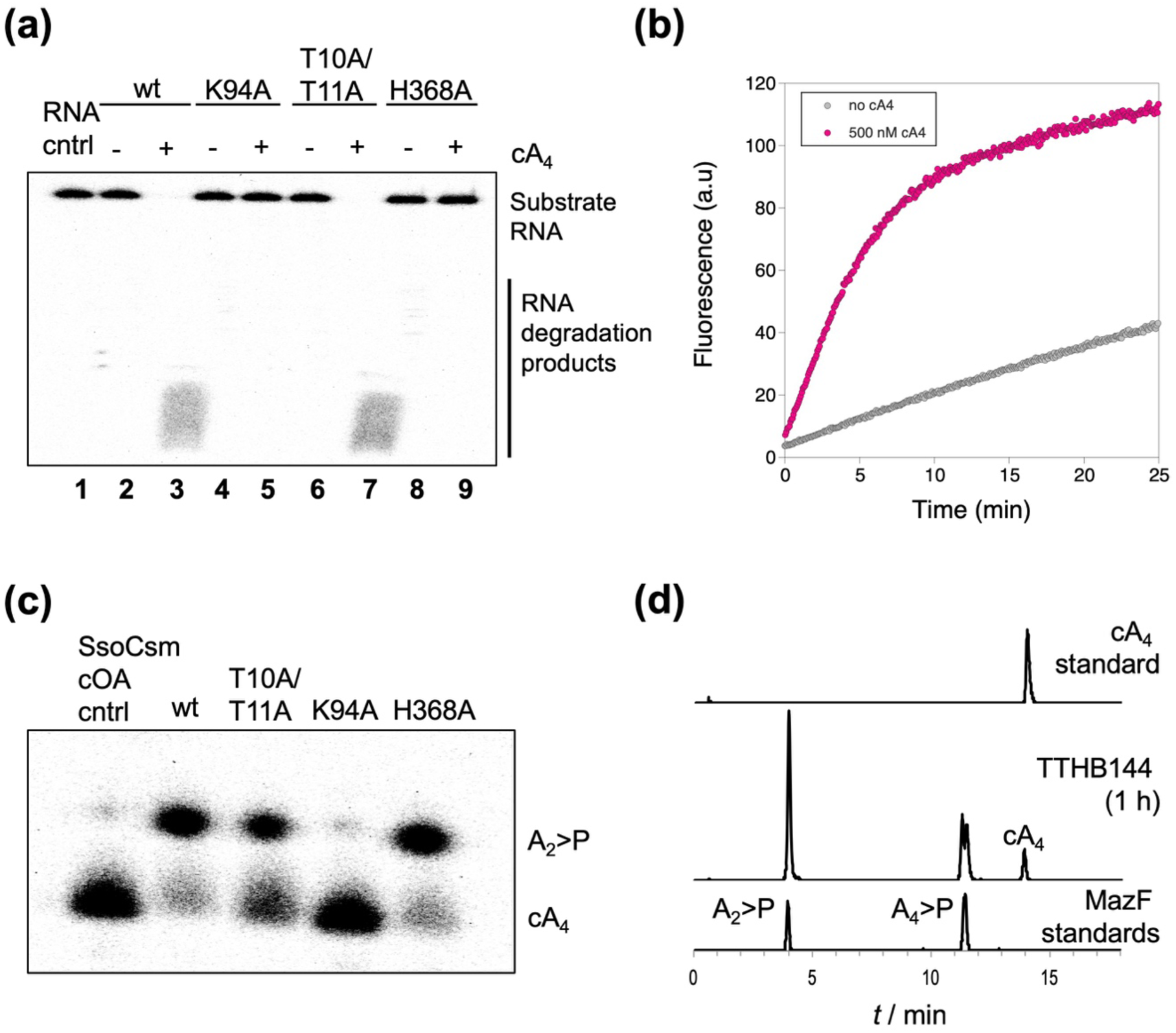
RNA and cA_4_ cleavage assays identify that cA_4_ is degraded at the CARF domain and cA_4_ is cleaved to di-adenylate containing a 5′ hydroxyl and a 2′,3′-cyclic phosphate. (**a**) Phosphorimage of denaturing polyacrylamide gel electrophoresis (PAGE) visualising the degradation of 50 nM radiolabelled A1 RNA, as previously described in [21], by TTHB144 (0.5 μM dimer) and its CARF domain variants K94A and T10A/T11A and the HEPN domain variant H368A. The reaction was incubated at 70 °C for 60 min in pH 8.0 buffer containing 20 mM Tris-HCl, 150 mM NaCl, 1 mM EDTA and 3 units SUPERase•In™ Inhibitor, before quenching by phenol-chloroform extraction. RNA was cleaved by wild-type (wt) protein and the T10A/T11A variant in the presence of 1 μM cyclic tetra-adenylate (cA_4_), but not by the K94A or H368A variants. Lanes are numbered for ease of identification. (**b**) Plot of fluorescence emitted when RNaseAlert™ substrate (1.5 μM, Integrated DNA Technologies) was degraded by 125 nM dimer TTHB144 in the absence (grey) or presence (magenta) of 500 nM cA_4_ at 50 °C. Fluorimetry was carried out in a Cary Eclipse Fluorescence Spectrophotometer (Agilent Technologies) with excitation and emission wavelengths set to 490 nm and 520 nm, respectively. (**c**) Phosphorimage of denaturing PAGE visualising degradation of 400 nM radiolabelled cA_4_ generated using *Sso*Csm complex, as previously described in [21], by TTHB144 (4 μM dimer) and variants at 70 °C for 120 min. cA_4_ was degraded to the slower migrating product di-adenylate containing a 5′ hydroxyl moiety and a 2′,3′-cyclic phosphate (A_2_>P). (**d**) High-resolution liquid chromatography mass spectrometry of cA_4_ produced using the *Sso*Csm complex and degradation products generated when the cA_4_ was incubated with TTHB144 (40 μM dimer) at 70 °C. cA_4_ (∼16 μM; top panel) was degraded to intermediate and product species (middle panel) with identical retention times to A_4_>P and A_2_>P, respectively (bottom panel). A_4_>P and A_2_>P standards were generated using the *E. coli* MazF toxin as previously described [8].

To investigate whether TTHB144 degraded cA_4_, we incubated the wild-type protein with radiolabelled cA_4_ generated using the *S. solfataricus* type III-D Csm complex [21]. TTHB144 degraded cA_4_ to generate a slower migrating product on denaturing PAGE (Figure 2c), which we have previously identified as di-adenylate containing a 5′ hydroxyl moiety and a 2′,3′-cyclic phosphate (A_2_>P) [13]. We verified this observation by high-resolution liquid-chromatography mass spectrometry, by comparison of cA_4_ degradation products to oligoadenylate standards generated using the *Escherichia coli* MazF toxin, as described previously [21]. Similar to the *S. solfataricus* ring nucleases, TTHB144 degraded cA_4_, yielding an A_4_>P intermediate and A_2_>P product (Figure 2d).

Subsequently, we evaluated cA_4_ degradation by TTHB144 CARF and HEPN domain variants. As *S. solfataricus* Csx1 does not degrade cA_4_ [13], we reasoned that cA_4_ degrading capability was unlikely to reside at the HEPN active site.. As anticipated, TTHB144 H368A, which has no ribonuclease activity, degraded cA_4_ similarly to wild-type protein, ruling out a role for the HEPN domain in cA_4_ degradation. However, cA_4_ degradation was abolished in the K94A variant and impaired in the T10A/T11A variant (Figure 2c), suggesting a role for the CARF domain in this reaction. To confirm this hypothesis, we quantified the single-turnover rates of cA_4_ degradation by TTHB144 and its active site variants at 70 °C. The wild-type and H368A variant degraded cA_4_ at rates of 0.012 ± 0.003 min^-1^ and 0.013 ± 0.002 min^-1^, respectively (Figure 3a&b) allowing us to definitively rule out the HEPN domain as the site of cA_4_ degradation.

**Figure 3.**
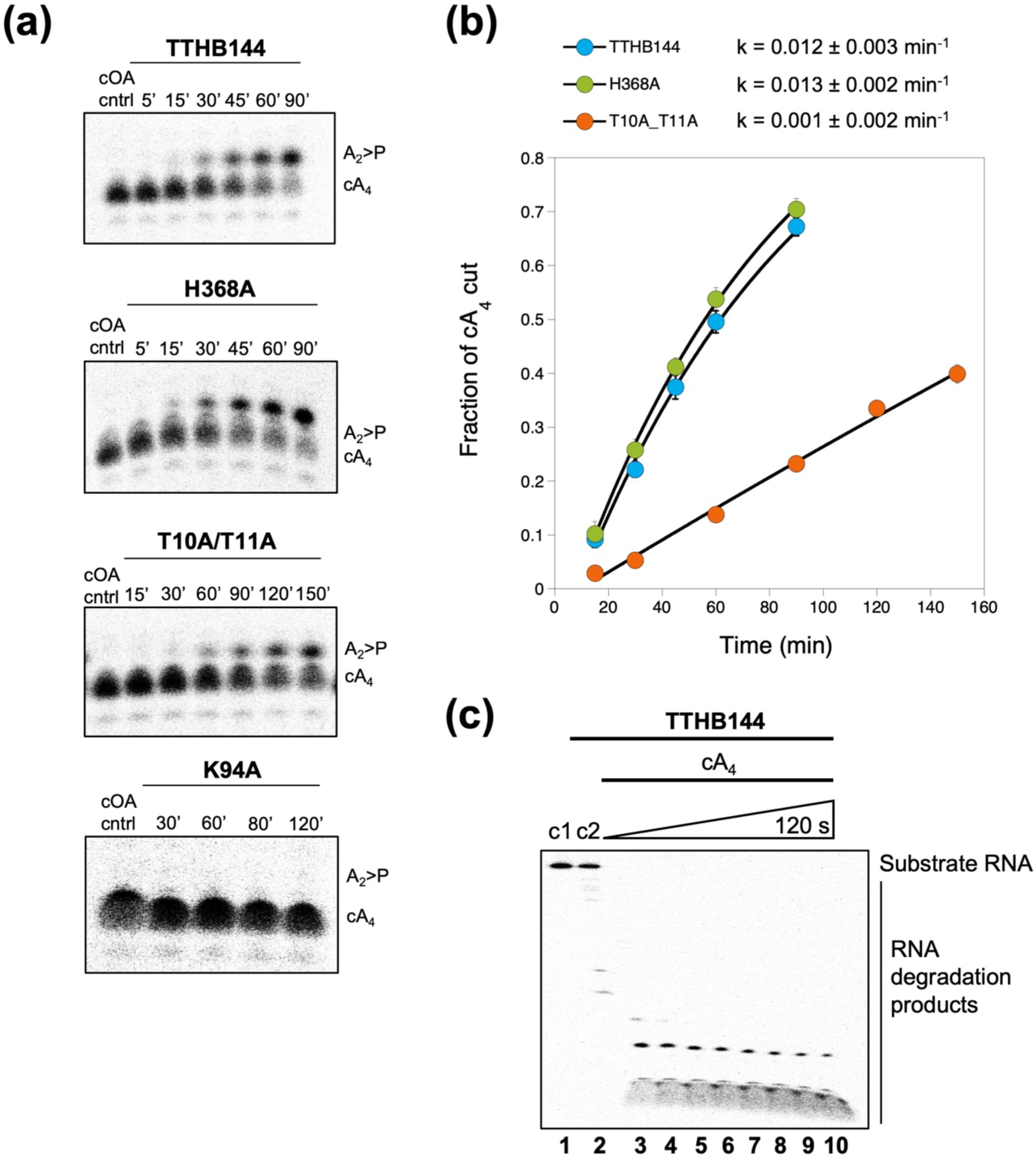
TTHB144 exhibits considerably slower kinetics of cA_4_ cleavage compared to RNA degradation under single turnover conditions. (**a**) Panels are phosphorimages of denaturing polyacrylamide gel electrophoresis (PAGE) visualising degradation of 200 nM radiolabelled cyclic tetra-adenylate (cA_4_) by TTHB144 and variants (8 μM dimer) at 70 °C. cA_4_ is degraded to A_2_>P. Time in minutes is indicated. Protein and radiolabelled cA_4_ were incubated in pH 8.0 buffer containing 20 mM Tris-HCl, 150 mM NaCl, 1 mM EDTA and 3 units SUPERase•In™ Inhibitor, and reactions were quenched at the indicated time-points by phenol-chloroform extraction. (**b**) Plot of the fraction of cA_4_ cut versus time, generated by quantifying the densiometric signals from the phosphorimages depicted in **a**. All data points are the average of three technical replicates and are fitted to an exponential rise equation to derive the rate of cA_4_ degradation, as described previously [21]. Error bars show the standard deviation of the mean. (**c**) Phosphorimage of denaturing PAGE visualising degradation of 50 nM radiolabelled RNA by TTHB144 (1 μM dimer) when incubated with 20 μM cA_4_ at 70 °C. Control reactions incubating RNA with buffer (c1) or RNA with protein in the absence of cA_4_ (c2) are shown. All of the substrate RNA was degraded within 15 s (lane 3).

The K94A variant was inactive, with no cA_4_ cleavage detectable over 2 h, consistent with a crucial role for this residue in cA_4_ binding, and the consequent loss of cA_4_-activated HEPN ribonuclease activity for this variant. In contrast, the T10A/T11A variant, which remains a cA_4_ activated HEPN ribonuclease (Figure 2a), exhibited a ∼12-fold decrease (0.001 ± 0.002 min^-1^) in cA_4_ cleavage compared to the wild-type protein. This suggests a role for these residues in cA_4_ degradation rather than binding. During cA_4_ cleavage, the hydroxyl group of the threonine residue likely facilitates an internal nucleophilic attack by the 2′ hydroxyl of the ribose sugar upon the 3′ scissile phosphate, as observed previously for the equivalent residue, S11, of Sso2081 [13]. The highly conserved lysine residue K94 appears crucial for cA_4_ binding, and may also play a role in stabilisation of the developing transition state during cA_4_ degradation. Hence the TTHB144 cA_4_ cleavage mechanism appears to be similar to that of the crenarchaeal ring nucleases [13]. Furthermore, to compare the rate of cA_4_ cleavage and cA_4_ stimulated RNA degradation, we sought to assay the rate of RNA degradation under single-turnover conditions. TTHB144 fully degraded the RNA within 15 s (Figure 3c), suggesting a lower limit of 10 min^-1^ for the catalytic rate constant. Thus, the rate of RNA degradation exceeds the rate of cA_4_ cleavage by approximately 3 orders of magnitude. It is worth noting that TTHB152, the previously characterised Csm6 from *T. thermophilus*, has a serine at position 11 and a lysine at the position equivalent to K94. Thus, TTHB152 may also catalyse degradation of cA_4_.

## Conclusions

TTHB144 is the first CARF family protein identified to harbour both cA_4_ degradation activity and ribonuclease activity. The single-turnover rate of cA_4_ degradation by TTHB144 at 70 °C is 2-fold lower than that of the least active *S. solfataricus* ring nuclease, Sso1393, at 60 °C. This slow rate of cA_4_ degradation may function as a built-in control mechanism to limit the extent of ribonuclease activity when in the activated state. *Streptococcus epidermidis* Csm6 activity during type III immunity has been shown to induce cell growth arrest [12], and self-limiting enzymes may be crucial for cell recovery in bacteria that do not have dedicated ring nucleases. Therefore, the amalgamation of a HEPN ribonuclease and a ring nuclease into a single self-limiting enzyme may help decrease the toxicity associated with non-specific RNA cleavage in type III CRISPR systems.

## Abbreviations

CARF: CRISPR Associated Rossman Fold
HEPN: Higher Eukaryotes and Prokaryotes Nucleotide binding
cOA: cyclic oligoadenylate
cA_4_: cyclic tetra-adenylate
TTHB: *Thermus thermophilus* HB8

## Acknowledgments

This work was funded by a grant from the Biotechnology and Biological Sciences Research Council (Grant REF BB/S000313/1 to MFW).

